# Characterization of RNA editing profiles in rice endosperm development

**DOI:** 10.1101/2024.01.27.577525

**Authors:** Ming Chen, Lin Xia, Xinyu Tan, Shenghan Gao, Sen Wang, Man Li, Yuansheng Zhang, Tianyi Xu, Yuanyuan Cheng, Yuan Chu, Songnian Hu, Shuangyang Wu, Zhang Zhang

**Author notes:** Corresponding authors: Zhang Zhang, Shuangyang Wu and Songnian Hu. Medical Center of Hematology, Xinqiao Hospital, State Key Laboratory of Trauma, Burn and Combined Injury, Army Medical University, Chongqing, China (Lin Xia). Beijing Key Laboratory for Agriculture Application and New Technique, College of Plant Science and Technology, Beijing University of Agriculture, Beijing, China (Sen Wang). Gregor Mendel Institute, Austrian Academy of Sciences, Vienna BioCenter, Dr. Bohr-Gasse 3, Vienna, Austria. These authors contributed equally to this work (Ming Chen, Lin Xia).

## Abstract

Rice (*Oryza sativa L*.) endosperm provides nutrients for seed germination and determines grain yield. RNA editing, a post-transcriptional modification essential for plant development, unfortunately, is not fully characterized during rice endosperm development. Here, we conduct genome re-sequencing and RNA sequencing for rice endosperms across five successive developmental stages and perform systematic analyses to characterize RNA editing profiles during rice endosperm development. We find that the majority of their editing sites are C-to-U CDS-recoding in mitochondria, leading to increased hydrophobic amino acids, and affecting structures and functions of mitochondrial proteins. Comparative analysis of RNA editing profiles across the five developmental stages reveals that CDS-recoding sites present higher editing frequencies with lower variabilities, and recoded amino acids, particularly caused by these sites with higher editing frequencies, tend to exhibit stronger evolutionary conservation across many land plants. Based on these results, we further classify mitochondrial genes into three groups that present distinct patterns in terms of editing frequency and variability of CDS-recoding sites. Besides, we identify a series of P- and PLS-class pentatricopeptide repeat (PPR) proteins with editing potential and construct PPR-RNA binding profiles, yielding candidate PPR editing factors related to rice endosperm development. Taken together, our findings provide valuable insights for deciphering fundamental mechanisms of rice endosperm development underlying RNA editing machinery.

**Author summary:** Rice endosperm development, a critical process determining quality and yield of our mankind’s essential food, is regulated by RNA editing that provokes RNA base alterations by protein factors. However, our understanding of this regulation is incomplete. Hence, we systematically characterize RNA editing profiles during rice endosperm development. We find that editing sites resulting in amino acid changes, called “CDS-recoding”, predominate in mitochondria, leading to increased hydrophobic amino acids and affecting structures and functions of proteins. Comparative analysis of RNA editing profiles during rice endosperm development reveals that CDS-recoding sites present higher editing frequencies with lower variabilities. Furthermore, evolutionary conservation of recoded amino acids caused by these CDS-recoding sites is positively correlated with editing frequency across many land plants. We classify mitochondrial genes into three groups that present distinct patterns in terms of editing frequency and variability of CDS-recoding sites, indicating different effects of these genes on rice endosperm development. In addition, we identify candidate protein factors associated closely with RNA editing regulation. To sum up, our findings provide valuable insights for fully understanding the role of RNA editing during rice endosperm development.

## Introduction

Rice (*Oryza sativa* L. subsp. *Geng* (*Japonica*) cv. Nipponbare) is a staple crop feeding more than half the world’s population [1] as well as a model plant for genetic studies and molecular breeding [2, 3]. The development of rice endosperm, rich in starch granules and protein bodies [4], determines grain production and quality [5]. Rice endosperm development is roughly divided into four stages: (i) coenocyte (1-2 days after flowering (DAF) or pollination; the pollination occurs at or slightly before flowering [6, 7]), (ii) cellularization (3-5 DAF), (iii) storage product accumulation (6-20/21 DAF), and (iv) maturation (21/22-30 DAF) [4, 8]. Among these stages, it is at stage III that accumulation of storage products and programmed cell death (PCD) simultaneously occur and play crucial roles in endosperm development [8, 9].

Plant mitochondria provide energy for endosperm development and are related to PCD [10, 11]. For rice, its mitochondrial genome contains 53 protein-coding genes, falling into four categories: (i) complexes composing the mitochondrial electron transport chain (mETC), including complex I (NADH dehydrogenase), complex III (cytochrome *c* reductase), complex IV (cytochrome *c* oxidase) and complex V (adenosine triphosphate (ATP) synthase), (ii) cytochrome (Cyt) *c* biogenesis proteins, (iii) ribosomal proteins, and (iv) other proteins [12]. Furthermore, mETC tightly couples with ATP synthesis; besides, it has been found that complexes I and III in the mETC produce reactive oxygen species (ROS) related to PCD [13, 14]. In mitochondria, two *c*-type cytochromes, namely, Cyt *c* and *c*_1_, participate in multiple biological processes; Cyt *c* transfers electrons from complex III to IV [15] and likely regulates PCD [16, 17], and Cyt *c*_1_ is integrated into complex III [15]. In addition, MatR, a maturase-related protein, is assigned to the “other proteins” group and is required for the splicing of Group II introns in mitochondria [18, 19]. Ribosomal proteins are responsible for translating mRNAs encoded by mitochondrial genomes and regulating protein synthesis, such as complexes in the mETC [20, 21].

RNA editing, as one of the most important post-transcriptional modifications, creates RNA products that differ from their DNA templates. RNA editing presents unique patterns of editing types in land plants, namely, mainly cytidine-to-uridine (C-to-U) alterations in mitochondria and plastids [22]. Importantly, pentatricopeptide repeat (PPR) proteins including P- and PLS-class type specifically bind RNA by PPR codes [23] and catalyze C-to-U RNA editing by DYW domain in PLS-class PPR proteins [24], playing a key role as editing factors [25]. Consequently, over the past decades, valuable efforts have been devoted to investigating RNA editing during plant endosperm development. Evidence has been accumulated that RNA editing in mitochondrial transcripts plays a key role in endosperm development [26–34] by affecting functions of complexes comprising mETC, Cyt *c* biogenesis proteins, ribosomal proteins and MatR. Specifically, RNA editing in mitochondrial genes regulated by PPR proteins is related to rice endosperm development, as *nad7* mediated by SMK1 [35], *atp6*, *cob*, *cox3*, *nad1*, *nad9*, *rpl16* and *rps12* by EMP5 [36], *cox2*, *cox3*, *ccmC*, *nad2* and *nad4* by OGR1 [37], and multiple mitochondrial genes (such as *nad9*) by cooperation of P-class FLO22 and PLS-class DYW3 [38]. As a result, RNA editing in mitochondrial genes as well as PPR editing factors have been identified in wild-type and mutant rice endosperm at specific developmental stages (for details see the Plant Editosome Database [39]), yet lacking a comprehensive characterization of RNA editing profiles during rice endosperm development.

Toward this end, here we collect rice endosperms across five successive developmental stages, mainly including the process of grain filling (stage III of rice endosperm development), and perform genome re-sequencing and strand-specific RNA sequencing (RNA-seq). Based on these sequencing data, we systematically profile RNA editing events, analyze RNA editing features in terms of location, editing frequency & variability, evolutionary conservation and structure, identify PPR genes in rice genome and construct PPR-RNA binding profiles, which collectively provide insightful clues for better understanding rice endosperm development from RNA editing machinery.

## Results

### RNA editing profiling during rice endosperm development

To systematically investigate RNA editing events during rice endosperm development, we obtain both genome re-sequencing and strand-specific RNA-seq data from rice endosperm derived from five successive developmental stages (3, 6, 9, 12, and 15 DAF). After strict quality control, sequence alignment, and filtration (see Materials and Methods), we identify a total of 369 high-confidence RNA editing sites across the five developmental stages (Table S1). Specifically, the primary RNA editing type in rice mitochondria is C-to-U (298) and the rest of types are G-to-A (65) and U-to-C (1), whereas in plastids the only type is C-to-U (5), revealing the dominance of C-to-U editing during rice endosperm development (Figure 1A and Table S2). For each developmental stage, the number of C-to-U editing sites in mitochondria is consistently larger in contrast to other editing types (Figure 1B). Furthermore, we capture a whole view of RNA editing events in rice mitochondrial genome across the five developmental stages (Figure 1C). We find that the average editing frequency of all 298 C-to-U sites in mitochondria varies from 2% to 99.8% at the five developmental stages, with the median at 69.8% (Figure 1D). Besides, the distribution of the distance between two neighboring editing sites shows that C-to-U editing sites tend to be located within 50bp (Figure S1). Among these 298 C-to-U editing sites in rice mitochondria, we observe that there are 253 sites located in coding sequence (CDS) regions (214 for CDS-recoding and 39 for CDS-synonymous), which are more prevalent by comparison against intronic (73), intergenic (12) and pseudogenic (8) regions (Figure 1E). Since one gene may be overlapped with other genes in the plant mitochondrial genome, notably, we find that among these 73 intronic editing sites, 43 are located in CDS (35 for CDS-recoding and 8 for CDS-synonymous) regions and 5 are in pseudogenic regions (Figure 1E and Table S1). Unlike the diversity of C-to-U editing distribution, the 65 G-to-A editing sites in mitochondria are located in intronic (23, 35.38%) and intergenic (42, 64.62%) regions and the 5 C-to-U editing sites in plastids are only in CDS regions (2 for CDS-recoding and 3 for CDS-synonymous) (Table S1). Taken together, it is clearly shown that CDS- recoding sites are more ubiquitous in mitochondria during rice endosperm development.

**Figure 1.**
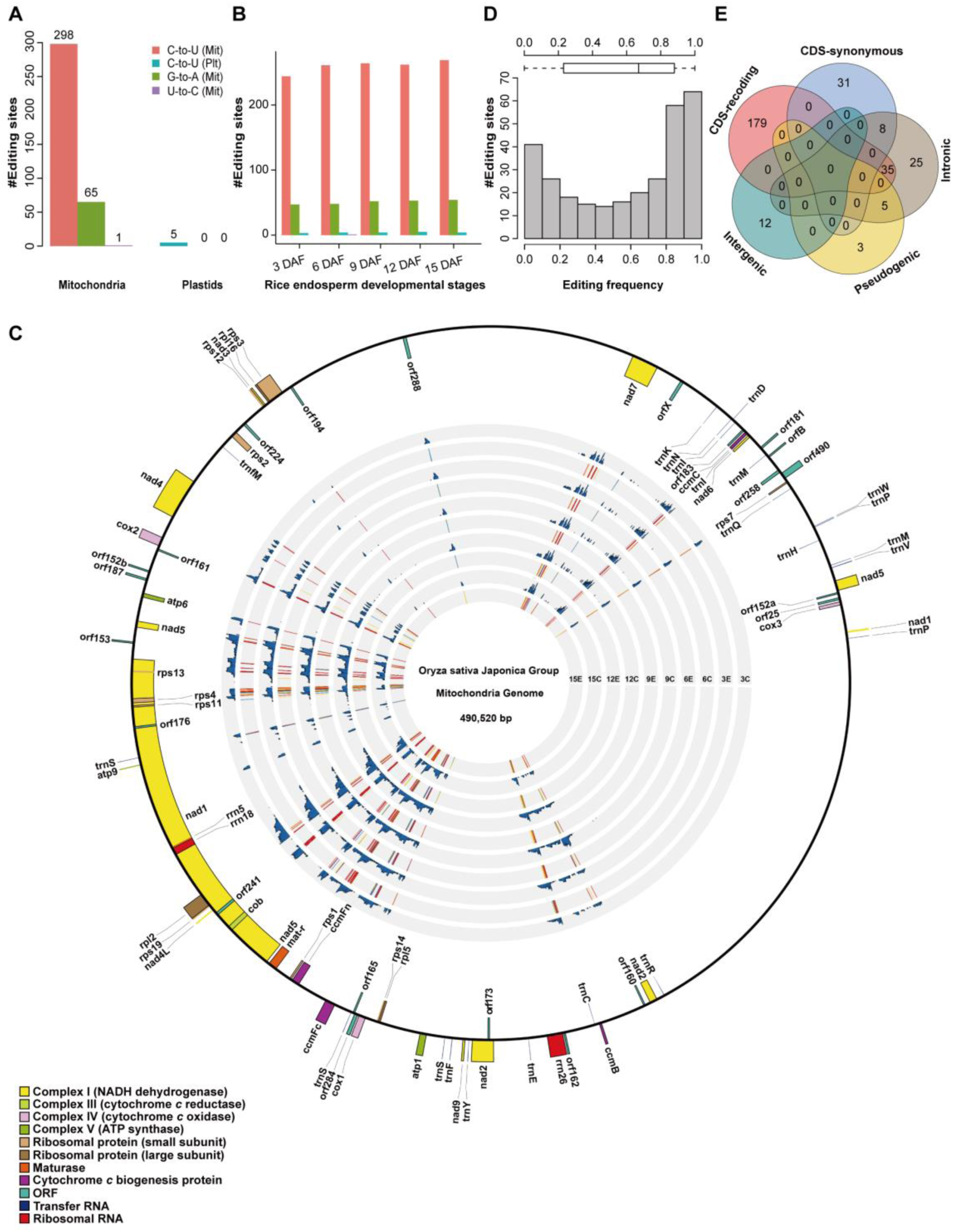
Characteristics of RNA editing profiles during rice endosperm development. **(A)** RNA editing sites in rice mitochondria and plastids. 369 RNA editing sites in mitochondria (364) and plastids (5) are categorized by editing types, which are color-coded, with pink for C-to-U (mitochondria), blue for C-to-U (plastids), green for G-to-A (mitochondria) and purple for U-to-C (mitochondria), respectively. **(B)** Distribution of 369 RNA editing sites at 3, 6, 9, 12, and 15 DAF. Bars with pink, green, blue, and purple denote C-to-U (mitochondria), G-to-A (mitochondria), C-to-U (plastids), and U-to-C (mitochondria) RNA editing types, respectively. **(C)** Distribution of RNA editing events and their corresponding RNA-seq coverage in rice mitochondria. The outer circle denotes rice mitochondrial genes; genes outside the circle are on the positive strand and oriented clockwise, while genes inside the circle are on the negative strand and oriented anticlockwise. The grey circles, from outer to inner denote RNA-seq coverage (Bar plot, C: Coverage) and frequency (Heatmap, E: Editing) of RNA editing sites at 3, 6, 9, 12, and 15 DAF. **(D)** Distribution of editing frequency of 298 C-to-U editing sites. **(E)** Distribution of 298 C-to-U editing sites in different sequence regions. Abbreviations used are: Mit, mitochondria; Plt, plastids.

### Amino acid biochemical properties changed by RNA editing

Studies have shown that RNA editing is functionally important by affecting amino acid biochemical properties [40–42]. In our study, among these 214 CDS- recoding sites, we find that 88 (41.12%) and 126 (58.88%) are located at the first and second codon positions, respectively. Contrastingly, among these 39 CDS-synonymous sites, most (36, 92.31%) occur at the third codon position producing plenty of U-ending codons, except 3 sites (7.69%; Leu-to-Leu, 2 for <u>C</u>UG-to-<u>U</u>UG and 1 for <u>C</u>UA-to-<u>U</u>UA) at the first codon position (Table S3 and S4). Accordingly, we identify 8 codon alteration types with more than 10 editing events: U<u>C</u>A-to-U<u>U</u>A (Ser-to-Leu), <u>C</u>GG-to-<u>U</u>GG (Arg-to-Trp), U<u>C</u>G-to-U<u>U</u>G (Ser-to-Leu), U<u>C</u>U-to-U<u>U</u>U (Ser-to-Phe), C<u>C</u>A-to-C<u>U</u>A (Pro-to-Leu), U<u>C</u>C-to-U<u>U</u>C (Ser-to-Phe), <u>C</u>GU-to-<u>U</u>GU (Arg-to-Cys), and UU<u>C</u>-to-UU<u>U</u> (Phe-to-Phe) (Figure 2A and Table S4). Considering the relative frequency of amino acids corresponding to these 253 sites in CDS regions (214 for CDS-recoding and 39 for CDS-synonymous) after RNA editing, Leu (94, 37.30%) and Phe (55, 21.83%) are relatively dominant (Figure 2A and Table S4). Regarding the amino acid biochemical properties, we find that nearly all amino acids after RNA editing on these 253 sites in mitochondria are hydrophobic except Ser and Thr (Figure 2A); the percentage of hydrophobic amino acids is dramatically increased from 37.70% to 86.90% by most CDS-recoding editing sites (188/214) (Table S4).

**Figure 2.**
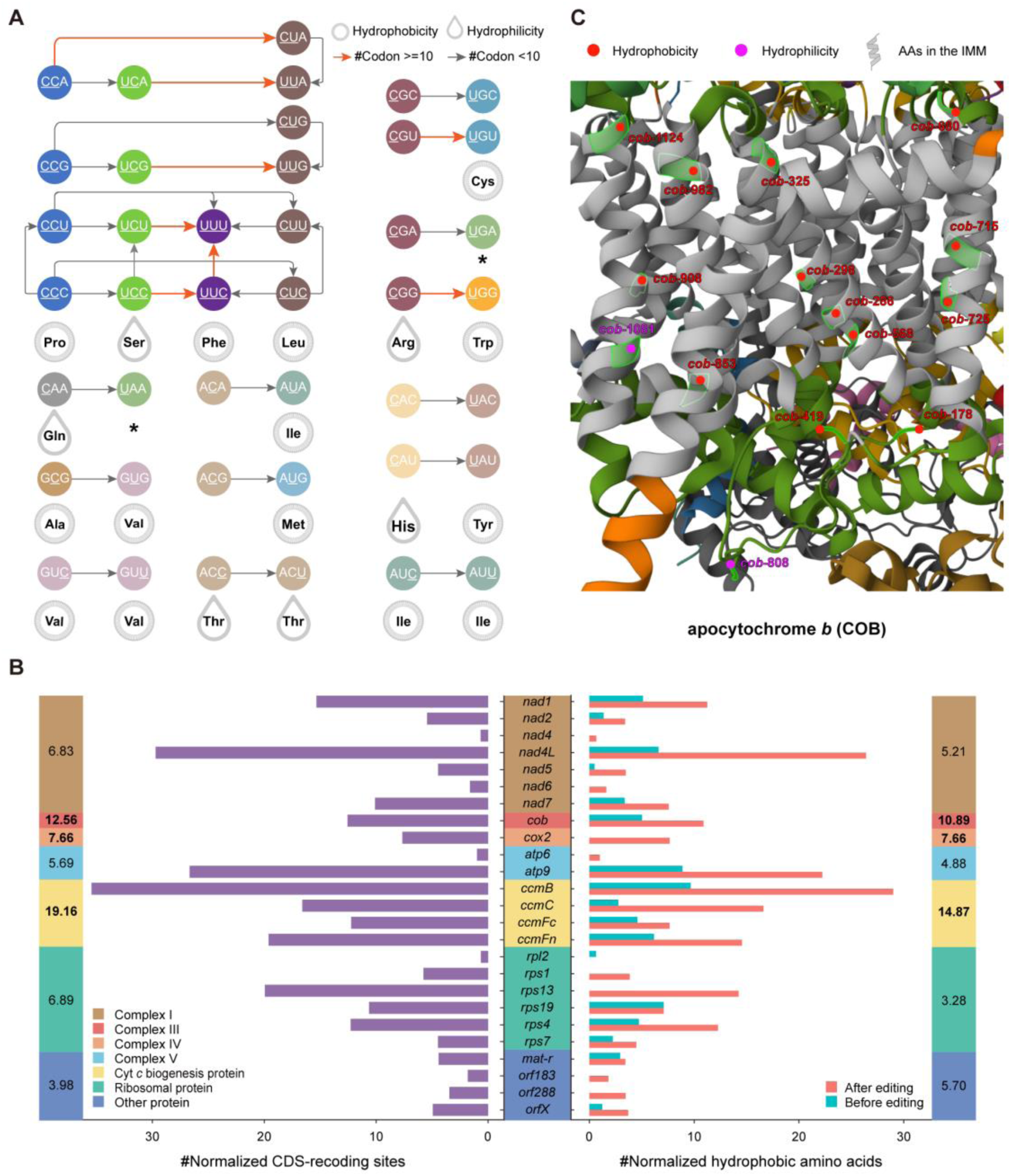
Codon alterations caused by editing in CDS regions, and the distributions of CDS-recoding sites and the corresponding recoded amino acids (AAs). **(A)** Amino acid and codon alterations of 253 editing sites in CDS regions. Red and grey arrows denote codon alterations with more and less than 10 editing events, respectively. Edited RNA bases are underlined in codons. Amino acids in membrane structures and water drops denote hydrophobic and hydrophilic amino acids, respectively. Asterisks denote stop codons. Abbreviations of amino acids are: Ala, Alanine; Arg, Arginine; Cys, Cysteine; Gln, Glutamine; His, Histidine; Ile, Isoleucine; Leu, Leucine; Met, Methionine; Phe, Phenylalanine; Pro, Proline; Ser, Serine; Thr, Threonine; Trp, Tryptophan; Tyr, Tyrosine; Val, Valine. **(B)** Normalized CDS-recoding site counts in different grouped genes as well as their corresponding change counts of amino acid hydrophobicity before and after editing. Purple bars denote normalized CDS-recoding site counts in protein-coding genes, equaling “CDS- recoding site counts / total length of CDS in the corresponding gene × 1000”. Numbers in the left and vertical bars denote total normalized CDS-recoding site counts in different types of proteins/complexes (complex I, III, IV, V, Cyt *c* biogenesis proteins, ribosomal proteins and other proteins from up to down). Normalized counts of hydrophobic amino acids corresponding to editing sites before and after editing in protein-coding genes are shown as blue and red bars, respectively. Normalized hydrophobic amino acid counts after editing in different types of proteins/complexes are shown in the right and vertical bars. In addition, calculation of normalized editing site counts in proteins/complexes, normalized counts of hydrophobic amino acid corresponding to editing sites before and after editing in protein-coding genes and proteins /complexes is similar to that of normalized editing site counts in protein-coding genes. **(C)** Recoded amino acids caused by RNA editing with mapped positions in the 3D structure of apocytochrome *b* (COB). The abbreviation used is: IMM, inner mitochondrial membrane.

In terms of 53 rice mitochondrial protein-coding genes assigned to 7 groups encoding complex I, complex III, complex IV, complex V, Cyt *c* biogenesis proteins, ribosomal proteins, and other proteins, we investigate the distribution of these 214 CDS-recoding sites as well as the corresponding amino acid hydrophobicity (Figure 2B). Among these 53 genes, nearly half (25 genes) harbor at least one CDS-recoding site (Figure 2B). Notably, 64 among these 214 CDS-recoding sites have been experimentally verified to be important for endosperm development in *Oryza sativa* [35, 37], *Zea mays* [27, 33, 34, 43] and *Arabidopsis thaliana* [34] (for details see Table S5), indicating the great potential of the rest CDS-recoding sites critical for endosperm development. Meanwhile, we detect RNA editing hotspots that feature more CDS-recoding sites and more hydrophobic amino acids after editing. We find that three gene groups, encoding Cyt *c* biogenesis proteins (normalized CDS-recoding sites: 19.16, normalized hydrophobic amino acids caused by CDS-recoding editing: 14.87), complex III (normalized CDS-recoding sites: 12.56, normalized hydrophobic amino acids caused by CDS-recoding editing: 10.89) and IV (normalized CDS-recoding sites: 7.66, normalized hydrophobic amino acids caused by CDS-recoding editing: 7.66) (Figure 2B), are RNA editing hotspots, indicating their key roles in rice endosperm development. In plastids, *atpA*-1148 is the only CDS-recoding site that presents higher editing frequencies (0.838 on average) (Table S1 and S6), which has been reported to be critical for ATP synthesis [44].

### Structural effects provoked by RNA editing

Considering the possible effect of RNA editing in protein structure [41, 45], we map these 214 CDS-recoding sites to three-dimensional (3D) structures of edited gene products. Notably, we observe that within mitochondria, most CDS- recoding sites (167/188) featuring hydrophobic amino acids are situated in/ near the inner membrane and/or the interior of proteins, characterized by a hydrophobic lipid environment (Figure S2 and Table S7), suggesting that editing at these sites plays a crucial role in preserving the insertion of membrane-proteins, ensuring protein stability, and subsequently influencing assemblies and activities of mitochondrial complexes/proteins, as previously elucidated [41]. Especially, *cytochrome b* (*cob*), encoding apocytochrome *b* (COB), a subunit of complex III in mitochondria that is one of RNA editing hotspots, possesses 15 CDS-recoding sites, among which 13 produce hydrophobic amino acids, conforming well with its hydrophobic environment (Figure 2C and Table S7). Likewise, in plastids, *atpA*-1148 and *ndhG*-347 causing hydrophobic amino acids are in/near IMM and/or buried in the interior of proteins (Figure S2 and Table S7).

To further investigate the effect of these CDS-recoding sites on protein 3D structure, we align 3D structures of mitochondrial proteins before and after editing. Obviously, we detect a significant difference (Root Mean Square Deviation, RMSD > 2.5Å according to the previous study [46]) in 3D structures of proteins encoded by *atp6*, *ccmB*, *ccmC*, *ccmFc*, *ccmFn*, *cob*, *cox2*, *mat-r*, *nad5*, *nad6*, *orf183*, *orf288*, *orfX*, *rpl2*, *rps1*, *rps4* and *rps19* (Figure S3), indicating that CDS-recoding editing in these genes greatly affects protein 3D structures. Notably, the RNA editing hotspots identified above, all present significant changes in 3D structures, especially *ccmB* (RMSD = 30.90Å) and *ccmFc* (RMSD = 25.06Å) (Figure S3). On the contrary, the rest mitochondrial protein-coding genes (*atp9*, *nad1*, *nad2*, *nad4*, *nad4L*, *nad7*, *rps7,* and *rps13*) delicately change their 3D structures with RMSD < 2.5Å (Figure S3), implying that CDS-recoding editing probably fine-tunes these proteins’ structures.

We also find that two intronic editing sites at *nad7*-i3-22 (position 22 at the 3^rd^ intron of *nad7*, genomic position at 87,186) and *ccmFc*-i-26 (position 26 at the only intron of *ccmFc*, genomic position at 332,291) affect the secondary structure of the domain 5 (D5) (Figure S4). The two RNA editing sites produce conserved GAAA in the apical loops of RNA secondary structure models of edited *nad7* intron 3 D5 and *ccmFc* intron D5 (Figure S4), suggesting their important functional roles in rice endosperm development. These results consist with previous findings that the conserved GAAA plays a key role in D5 functions [47, 48] and RNA editing at *nad7*-i3-22 affects splicing of *nad7*, production of NAD7 and maize endosperm development [49]. In addition, the effect of editing at *ccmFc*-i-26 on rice endosperm development needs further experimental verifications.

### RNA editing frequency, variability and evolutionary conservation

We next investigate the editing frequency and variability of these 298 C-to-U sites in mitochondria across the five developmental stages. For each stage, it is consistently observed that CDS-recoding sites present significantly higher editing frequencies than CDS-synonymous, intronic and intergenic ones (Figure 3A). Strikingly, all CDS-recoding and CDS-synonymous sites in *cob*, encoding a subunit of complex III and belonging to the editing hotspot, feature higher editing frequencies at each developmental stage (Figure S5). As one site may present variable editing frequencies at different developmental stages, we further examine the editing variability of these sites. We find that CDS- recoding sites also possess lower editing variability than other sites (Figure 3B), indicating the significant role of their relative constant editing in rice endosperm development. Specifically, we notice that most CDS-recoding sites in *atp6*, *ccmC*, *cox2*, *nad4L*, *nad6*, *orf183* as well as ribosomal protein genes, present not only higher editing frequencies but also lower editing variabilities across the five developmental stages, clearly showing the critical significance of RNA editing in altering amino acids to yield functional proteins for rice endosperm development.

**Figure 3.**
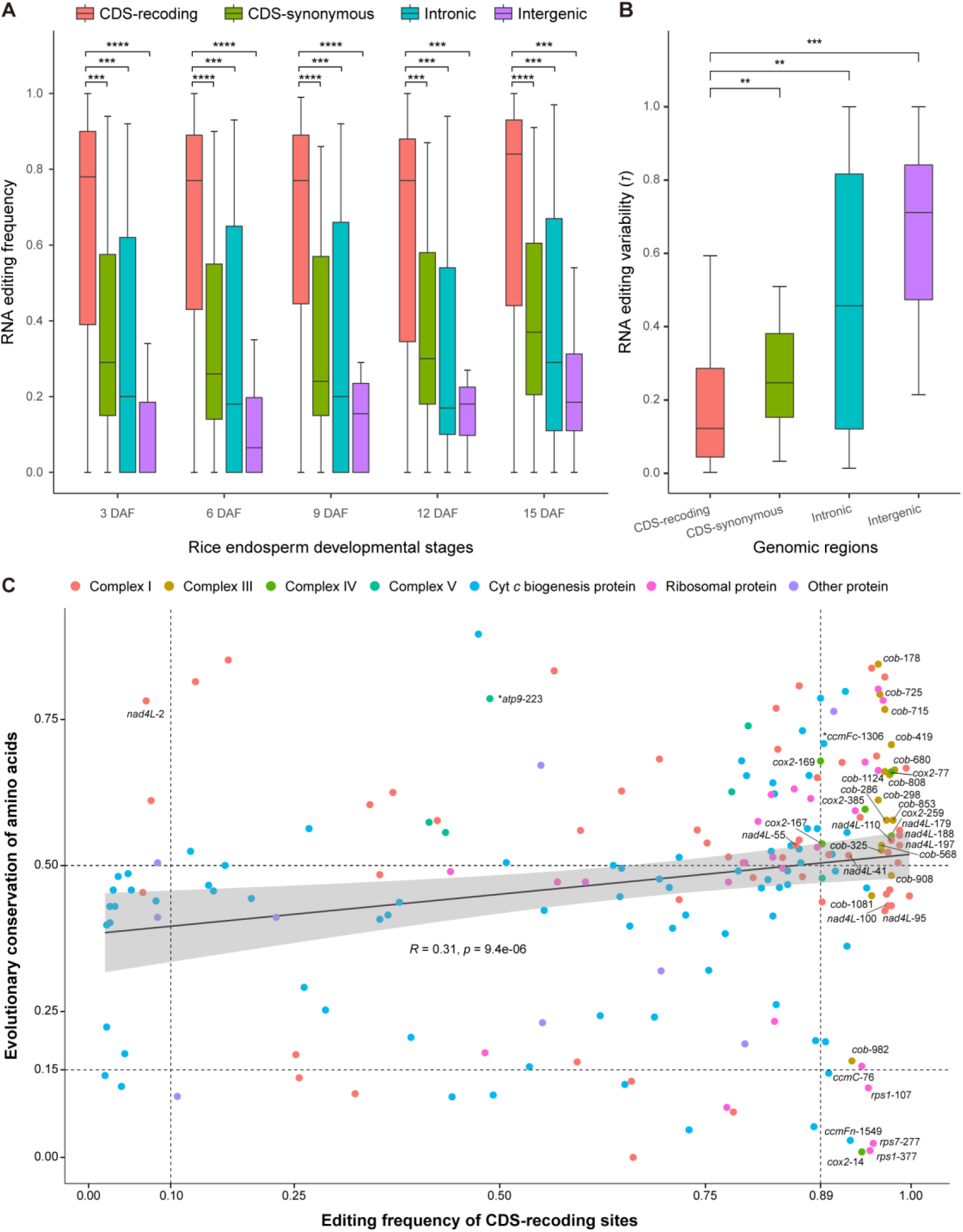
Developmental and evolutionary features of CDS-recoding events in mitochondria. **(A)** Editing frequencies of 247 C-to-U sites beyond overlapped genes in CDS (179 for CDS-recoding and 31 for CDS-synonymous), intronic (25) and intergenic (12) regions during five rice endosperm development stages. In addition, editing frequency of three sites in pseudogenic regions are not compared with that of CDS-recoding sites because of few sites and lack of sufficient statistical significance. The Wilcoxon test was used to calculate the statistical significance *p*-value, with ***<0.001, and ****<0.0001. **(B)** Editing variabilities of these 247 C-to-U sites across five stages during rice endosperm development. The Wilcoxon test was used to calculate the statistical significance *p*-value, with **<0.01, and ***<0.001. **(C)** Correlation between editing frequency of CDS-recoding sites and evolutionary conservation of amino acids caused by these CDS-recoding sites. Points in different colors denote CDS-recoding sites in mitochondrial genes encoding different proteins/complexes. Based on the density distribution of conservation of amino acids (Figure S6), thresholds of high and low conservation were set at 0.50 and 0.15, respectively. Similarly, thresholds of high and low editing frequencies were set at 0.89 and 0.10, respectively, according to the density distribution of editing frequencies of these CDS-recoding sites (Figure S7).

RNA editing is believed to restore evolutionarily conserved amino acids [50]. In other words, for one CDS-recoding site, its recoded amino acid via RNA editing may be restored to the amino acid that is relatively conserved in multiple species. To explore such editing conservation of these 214 CDS-recoding sites, we retrieve a large collection of protein sequences corresponding to these 214 sites from 122 land plants, falling into 6 different clades, namely, liverworts (12), mosses (25), hornworts (5), lycophytes (3), ferns (2) and seed plants (75) (Table S8). Accordingly, we find that these CDS-recoding sites present a positive correlation between amino acid conservation and editing frequency (*R* = 0.31, *p* = 9.4e-06) (Figure 3C; for details see Figure S8), which, albeit the correlation coefficient is not so strong, is observed in nearly all clades except hornworts and ferns (Figure S9).

We further classify these editing sites into 7 groups in terms of gene products as mentioned above. We find that editing sites in genes encoding Cyt *c* biogenesis proteins exhibit the positive correlation between amino acid conservation and editing frequency (Figure S10). Particularly, *cob* (encoding a subunit of complex III), *cox2* (encoding a subunit of complex IV) and *nad4L* (encoding a subunit of complex I) tend to possess more conserved amino acids and higher editing frequencies (Figure 3C). On the contrary, it is also believed that RNA editing enriches proteomic diversity [51]. Notably, *ccmC*, *ccmFn*, *cox2*, *rps1*, and *rps7* harbor a few CDS-recoding sites with higher editing frequencies but less conserved amino acids, indicating the enriched proteomic diversity in these genes (Figure 3C).

It is worth noting that RNA editing can provoke initiation and stop codons. Here we find that these CDS-recoding editing events lead to one initiation codon (*nad4L*-2, A<u>C</u>G to A<u>U</u>G) and three stop codons (*atp9*-223, <u>C</u>GA to <u>U</u>GA; *ccmFc*-1306, <u>C</u>GA to <u>U</u>GA; and *rps1*-505, <u>C</u>AA to <u>U</u>AA) (Table S6). Intriguingly, *nad4L*-2 has lower editing frequency in rice but higher conservation in its encoded amino acid across 117 land plants (Figure 3C and Table S9), implying that during rice endosperm development ACG mainly acts as the initiation codon in *nad4L*, consistent with a previous finding in *Phaseolus vulgaris* mitochondria [52]. In addition, *atp9*-223 and *ccmFc*-1306 present higher conservation across land plants, leading to stop codons at the end of corresponding gene products (Figure 3C; Table S6 and S9). Contrastingly, RNA editing at *rps1*-505 resulting in a stop codon is observed only in rice and absent in other land plants, accordingly reducing the length of *rps1* products by 4 amino acids.

### RNA editing pattern and classification of mitochondrial genes

Since CDS-recoding editing sites predominate, and present important effects on biochemical properties and protein structures during rice endosperm development, we further investigate their editing patterns during the five developmental stages. Based on unsupervised clustering, we group the 214 CDS-recoding sites into five clusters, namely, C1∼C5, with each cluster presenting a distinctive RNA editing pattern, containing 26, 17, 25, 43, and 103 sites, respectively (Figure 4A). We find that the average editing frequency is on the increase from C1 to C5 (with 0.08 in C1, 0.28 in C2, 0.50 in C3, 0.74 in C4, and 0.92 in C5, respectively), and their corresponding editing variability is gradually on the decrease (with τ value at 0.85, 0.44, 0.25, 0.15 and 0.03, respectively; lower τ value indicates more relatively constant editing frequencies). Clearly, C5 features higher editing frequency and lower editing variability, whereas C1 features lower editing frequency and higher editing variability.

**Figure 4.**
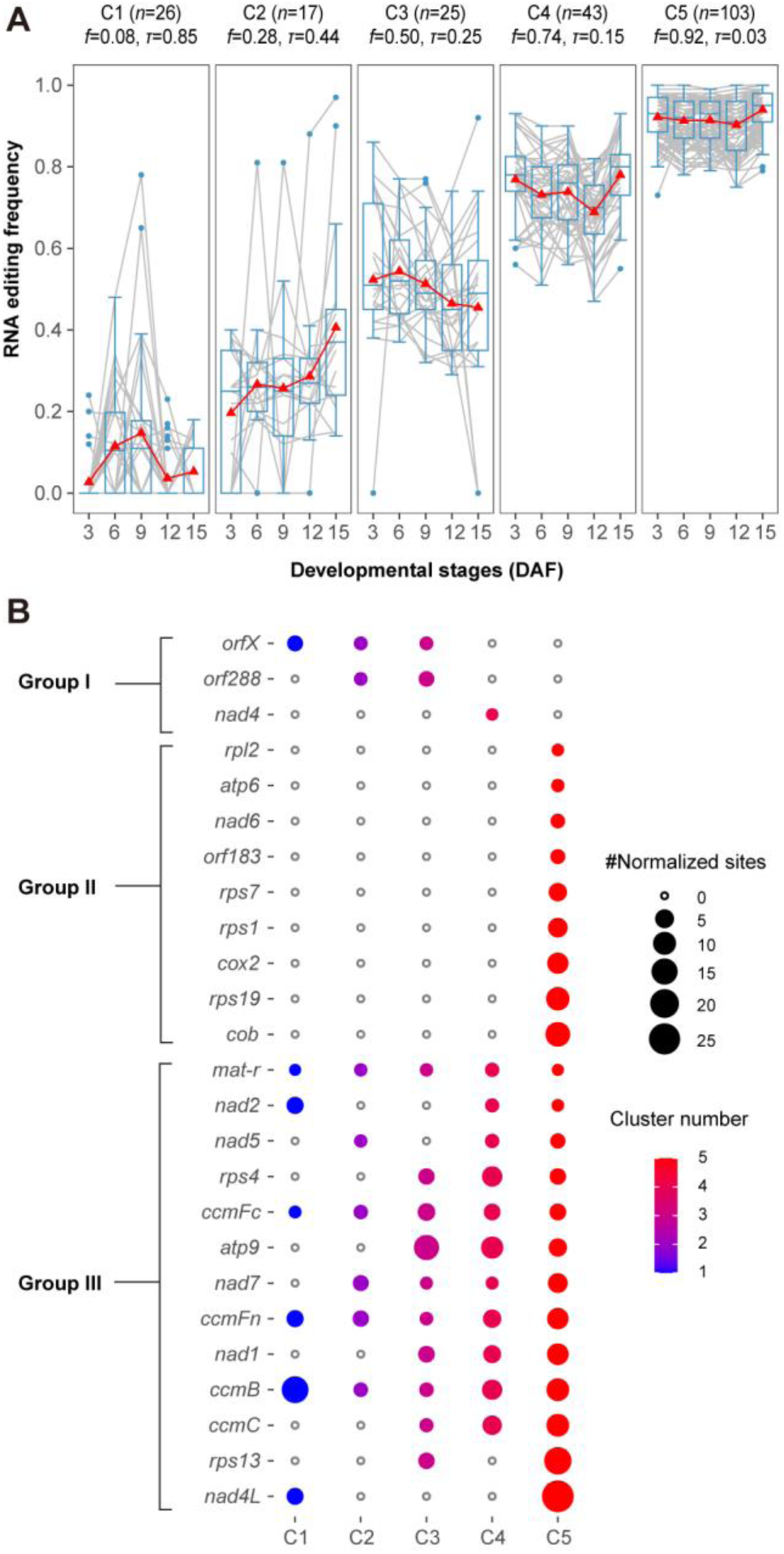
Classification of mitochondrial genes by editing frequency of 214 CDS-recoding sites. **(A)** Clustering of 214 CDS-recoding sites in five developmental stages of rice endosperm, leading to five clusters (C1∼C5) that show distinct patterns by factoring both editing frequency (“*f*”) and variability (“*τ*”). In each cluster, the red triangle denotes the average editing frequency in each development stage. **(B)** Classification of edited mitochondrial protein-coding genes based on the clustering patterns of these 214 CDS-recoding sites.

To further investigate 25 mitochondrial genes containing CDS-recoding sites, we classify these genes, by factoring C1∼C5, into Groups I, II, and III which contain 3, 9, and 13 genes, respectively. Specifically, Group I harbors editing events present in C1∼C4 yet absent in C5, Group II features C5-specific editing events, and Group III contains editing events spanning broadly from C1∼C5 (Figure 4B). In particular, in Group I, *nad4* encoding a subunit of complex I contains CDS-recoding events only from C4. Of note, *cob* encoding a component of complex III and *cox2* encoding a component of complex IV, belonging to Group II, are RNA editing hotspots. Similarly, four genes (*ccmB*, *ccmC*, *ccmFc* and *ccmFn*) encoding Cyt *c* biogenesis proteins in Group III are also RNA editing hotspots, with diverse CDS-recoding events covering all five clusters (C1∼C5) except *ccmC*. Furthermore, since editing frequencies are increased or decreased in mutant endosperms (Table S5), regulating RNA editing in genes from Groups I and III is potential to affect functions of mitochondrial complexes/proteins and further improve rice endosperm development.

### Construction of PPR-RNA binding profiles

PPR proteins are representative RNA editing factors that specifically bind RNA by PPR codes and directly modulate RNA editing by DYW domain [24]. To explore PPR proteins regulating RNA editing during rice endosperm development according to the distinctive mechanism of PPR-RNA binding, we identify 480 PPR genes in rice genome, among which half are PLS-class (including PLS-, E+-, E1-, E2-, and DYW-subtype) PPR genes and half P-class PPR genes (Figure 5A and Table S10). To further predict potential RNA editing factors (for details see Materials and Methods), we find that a total of 197 P-class and 225 PLS-class PPR proteins are likely to participate in RNA editing in mitochondria and/or plastids (Table S11). Furthermore, based on PPR codes and their binding affinities for RNA, we predict PPR-RNA binding and construct P- and PLS-class PPR-RNA binding profiles (Tables S12-S14). Specifically, we find that LOC_Os12g17080 and LOC_Os11g10740 tend to bind upstream RNA sequence containing the top 10 editing sites (Table S12). Among these top 10 sites, notably, *cox2*-167 (ranking 3^rd^) (Figure 5B) and *nad7*-836 (ranking 6^th^) (Figure 5C) have been experimentally validated to affect endosperm development in rice [35, 37], implying that our predicted PPR-RNA binding profiles provide valuable candidates for studying RNA editing machinery during rice endosperm development. Furthermore, we investigate the relationship between PPR gene expression and RNA editing frequency and find that PPR genes are lowly expressed during rice endosperm development (Table S15) conforming with previous findings [53–55], implying that PPR proteins are highly efficient editing factors. Moreover, the low expression of PPR detected above during rice endosperm development indicates that expression is not a good indicator for screening RNA editing factor candidates.

**Figure 5.**
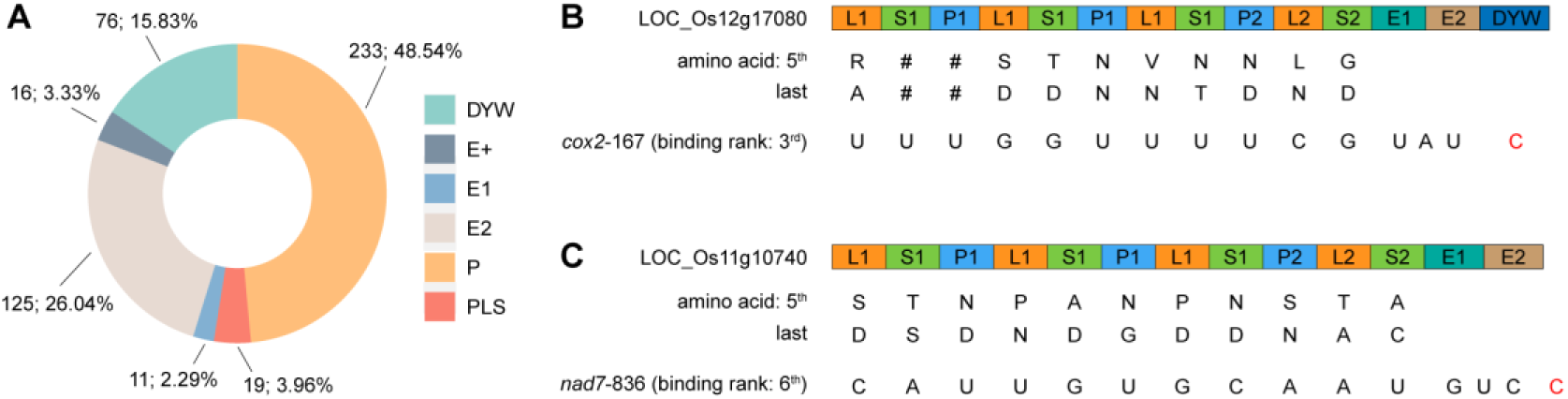
Identification of PPR genes and prediction of PPR-RNA binding. **(A)** A total of 480 PPR genes were identified in the rice genome, containing P- and PLS-class (including PLS-, E+-, E1-, E2-, and DYW-subtype according to compositions of different motifs) PPR genes. **(B and C)** Motif arrangement of LOC_Os12g17080 and LOC_Os11g10740 and their binding to the upstream sequence of *cox2*-167 and *nad7*-836, respectively. The combination of the fifth and last amino acids in each motif (P1, L1, S1, P2, L2 and S2) denotes a PPR code, specifically binding to a RNA base. The cytidine (C) in red is the edited RNA base.

## Discussion

Here, we constructed RNA editing profiles in mitochondria during rice endosperm development and performed systematic analyses to characterize RNA editing sites, demonstrating that CDS-recoding editing is conserved in evolution and development, increases the proportion of hydrophobic amino acids located in/near the mitochondrial inner membrane and/or interior of proteins, and changes 3D structures of mitochondrial proteins. Moreover, we classified mitochondrial protein-coding genes containing CDS-recoding editing events into three groups based on editing frequency and variability, and constructed PPR-RNA binding profiles to yield a high-confidence collection of candidate PPR editing factors. Based on these results, we present a model of RNA editing in mitochondria for rice endosperm development (Figure 6): RNA editing modulated by one PPR protein changes RNA and amino acid sequences and affects complex/protein formation to maintain ATP synthesis, protein synthesis, and PCD involving genes that encode complexes (complex I, III, IV and V) in the mETC, Cyt *c* biogenesis proteins and ribosomal proteins, which together sustain rice endosperm development.

**Figure 6.**
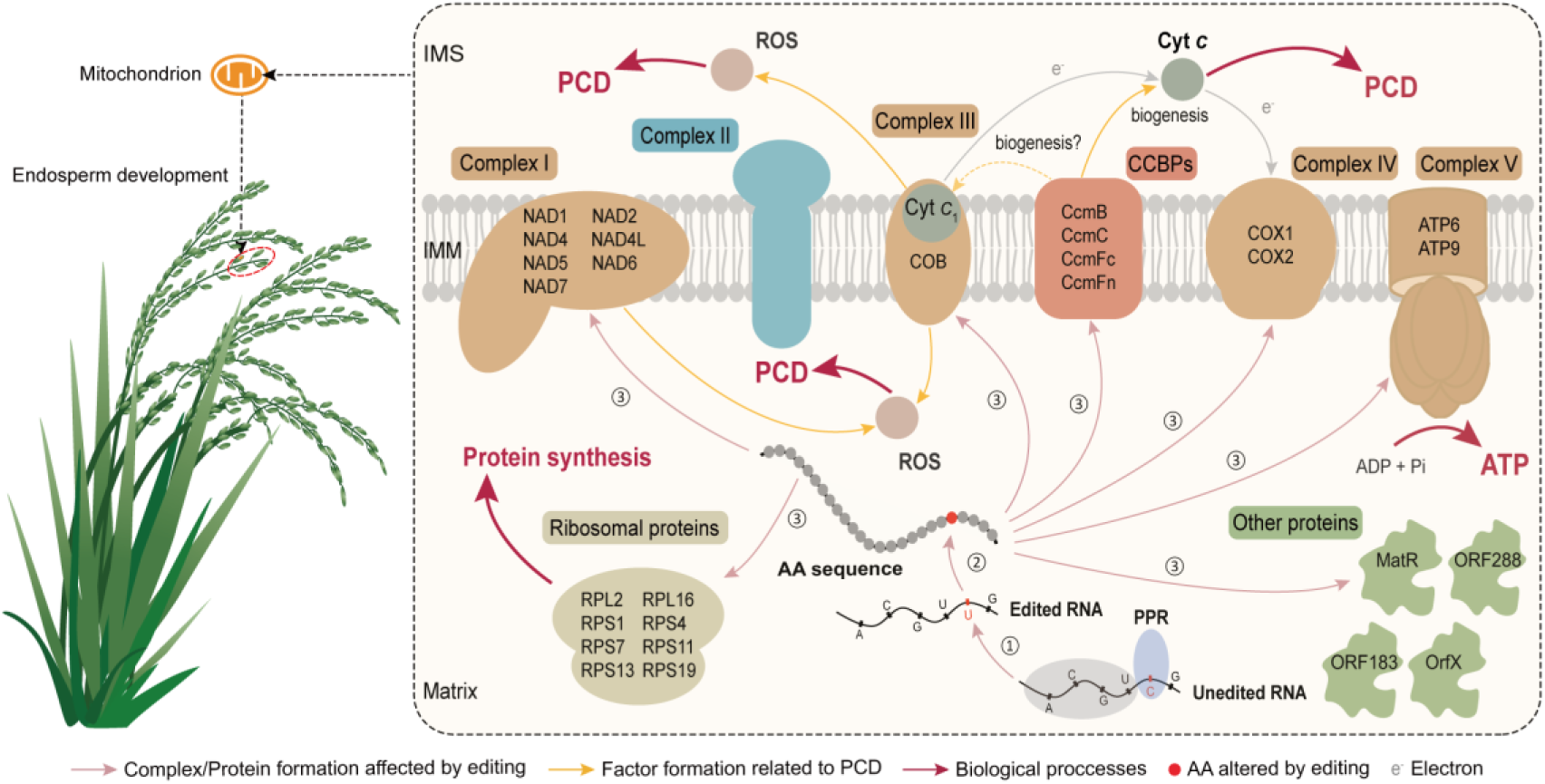
A model of CDS-recoding editing in mitochondria for rice endosperm development. Editing regulated by one PPR protein changes RNA and amino acid sequences, regulates formations and functions of complexes (complex I, III, IV and V), Cyt *c* biogenesis proteins and ribosomal proteins to ensure ATP synthesis, PCD and protein synthesis, which together maintain rice endosperm development. Different gene categories containing CDS-recoding editing events are color-coded, with complexes (complex I, III, IV and V) for brown, Cyt *c* biogenesis proteins for pink, ribosomal proteins for grey, and other proteins for green. The yellow dashed line denotes CcmB, CcmFc and CcmFn are related to Cyt *c*_1_ biogenesis, while the question mark means that the role of CcmC in Cyt *c*_1_ biogenesis is uncertain. Complex II is encoded by two nuclear genes and is not found to contain RNA editing events in rice. Abbreviations used are: AA, amino acid; IMS, intermembrane space; IMM, inner mitochondrial membrane; CCBPs, cytochrome *c* biogenesis proteins; ROS, reactive oxygen species; Cyt, cytochrome.

CDS-recoding editing in genes encoding complexes (complex I, III, IV, and V) in the mETC and encoding Cyt *c* biogenesis proteins, plays a key role in ATP synthesis during rice endosperm development. For instance, *cox2*-167, an editing site identified in the gene encoding complex IV that is the last electron acceptor in the mETC and crucial for ATP formation, has been documented to present aberrant RNA editing leading to abnormal rice endosperm [37]. Strikingly, there is a body of related evidence supported by several studies on maize endosperm development: (i) absence of editing at *cob*-908 (also identified in this study) results in deficiency of complex III, decrease of ATP production and reduction of maize endosperm growth [27], (ii) loss of editing at *ccmB*-43 (also identified in this study) affects the maturation of Cyt *c* and *c*_1_, and biogenesis of complex III [34], and (iii) abolishment of editing at *ccmFn*- 1553 in maize (corresponding to *ccmFn*-1514 identified in this study) results in the loss of CcmFN protein (a subunit of heme lyase complex) and affects cytochrome *c* maturation, mitochondrial oxidative phosphorylation [33], which, as a result, affects maize endosperm development.

In addition, attention should be paid to CDS-recoding editing in genes that encode other mETC complexes, such as complex I and V, because, besides ATP synthesis, complex I is related to formation of ROS that is the crucial signaling molecule regulating PCD [13, 14]. Consistently, abolishment of editing at *nad2*-26 and *nad5*-1916 (also identified in this study) affects PCD and maize endosperm development [26]. Moreover, CDS-recoding editing events in *nad4L* (*nad4L*-110 and *nad4L*-179) encoding a subunit of complex I and *atp9* (*atp9*- 191 and *atp9*-212) encoding a subunit of complex V (also identified in this study), are closely associated with mutant maize endosperm [29, 43]. In addition, abolishment of editing at *nad7*-836 affects the structural stability and activity of NAD7 (a subunit of complex I) and further influences respiration rate, starch biosynthesis, and maize endosperm development [35].

CDS-recoding editing in ribosomal proteins is important for protein synthesis during rice endosperm development. Particularly, it has been reported that editing at *rps4*-335 (in maize, *A. thaliana,* and rice in this study), converting Pro into Leu, is related to the maturation of Cyt *c*, the assembly of complex I, III, and V, and affects protein translation and endosperm development [34]. In addition, ribosomal protein S1, S7 and S13 are essential to protein synthesis [56–58] and CDS-recoding editing events in *rps1* (*rps1*-107 and *rps1*-377), *rps7* (*rps7*-277 and *rps7*-332) and *rps13* (*rps13*-56, *rps13*-100, *rps13*-256 and *rps13*-287) (also identified in this study) affect mutant maize endosperm [32, 33, 43, 59, 60], implying that CDS-recoding in these genes plays a key role in protein synthesis during rice endosperm development.

CDS-recoding editing in genes encoding other mitochondrial proteins is also critical for rice endosperm development. Notably, 3D structures of genes *mat-r*, *orf183*, *orf288* and *orfX* are significantly changed after RNA editing, especially for *orf288* (RMSD = 24.57Å) and *mat-r* (RMSD = 16.77Å) (Figure S3), implying the importance of editing in these genes for rice endosperm development. Specifically, it has been reported that abnormal editing at *mat-r*-86, *mat-r*-92, *mat-r*-295, and *mat-r*-296 is potentially associated with mutant endosperm of maize (also identified in this study) [28, 43]. In addition, it has been found that abolishment of editing at one site in *mat-r* can reduce the splicing efficiency of *nad4* Introns 1 and 3 and further affect maize endosperm development [31]. However, *orfX*, *orf183,* and *orf288*, similar to a chimeric cytoplasmic male sterility-associated gene *orf312* [61], have not been well studied, requiring further investigations on uncovering molecular mechanisms and functions of RNA editing in these genes.

Meanwhile, CDS-synonymous editing, particularly with high editing frequency, can also affect rice endosperm development. Although, as known to all, CDS- synonymous editing events do not cause amino acid alterations, some of them, such as *cob*-1119, *cob*-564, and *ccmFn*-372, present relatively higher editing frequencies (Table S1), suggesting that synonymous RNA editing does play an important role in rice endosperm development. As previously reported, synonymous mutations created by RNA editing likely stabilize RNA molecules [62] and affect translation efficiency based on changing synonymous codon types [40].

## Materials and Methods

### Whole genome re-sequencing and RNA sequencing

DNA was extracted from 3 days after flowering (DAF) leaves and endosperm of the wild type *Nipponbare (Japonica)* by the DNA extraction kit (DP305) from HUAYUEYANG Biotechnology, BEIJING CO.,LTD. RNA was extracted from 3, 6, 9, 12, and 15 DAF by the RNA extraction kit (GK0416) produced by HUAYUEYANG Biotechnology, BEIJING CO.,LTD. Sequencing libraries for each sample were prepared parallelly by both the KAPA Stranded mRNA-seq Library Preparation kit (KK8421) and the Epicentre Ribo-Zero Magnetic Kit (Plant Seed/Root, MRZSR116). Both DNA and RNA raw sequencing data were generated by Illumina HiSeq 2500 instruments, and these data were deposited in the Genome Sequence Archive (GSA) [63] at CRA008750 in the National Genomics Data Center, part of Beijing Institute of Genomics, Chinese Academy of Sciences & China National Center for Bioinformation [64].

### Read mapping and gene expression profiling

Transcript assembly and quantification based on strand-specific RNA-seq data involved three steps: quality control, transcript assembly, and gene expression estimation. In brief, Trimmomatic (version 0.35) [65] was used to eliminate low-quality reads as well as to remove adapters. Next, cleaned reads were mapped to a complete rice genome including the mitochondria genome (NC_011033.1) [66], the chloroplast genome (NC_001320.1) [67] and the nuclear genome (Os-Nipponbare-Reference-IRGSP-1.0) [68] by HISAT2 (version 2.2.1) [69] with a customized parameter “*--rna-strandness FR*” to specify strand-specific information. Subsequently, the mapped reads were used to quantify expression levels with FPKM (Fragments Per Kilobase of exon per Million fragments mapped) for both genes and transcripts by stringTie [70].

### Identification of RNA editing sites and estimation of editing frequency

To detect editing sites, the matched RNA and genomic DNA samples were analyzed by REDItoolDnaRna.py program provided in REDItools [71]. In general, the key procedure of RNA editing identification is to detect the mismatches between DNA and RNA sequence. However, the DNA-RNA mismatches could occur due to Single Nucleotide Polymorphisms (SNPs), misalignments and sequencing errors. In order to identity high-quality RNA editing sites, SNPs were identified by comparing the re-sequenced DNA sequence to the reference genome and were excluded. To avoid the misalignments caused by gene transfers and duplicated regions, “*-e -E -d -D*” was used to remove multiple mapped reads as well as duplicated reads of both RNA and DNA sequencing data, while the reads mapped to mitochondria or chloroplast genomes were kept. In addition, to exclude the sequencing errors, low-quality bases (quality score <25) and low-mapping sites (quality score <25) were removed. DNA-RNA mismatches undergoing multiple changes were excluded, and only those supported by both RNA and DNA sequencing with base coverages greater than 30 and variations supported by at least 3 reads were kept. RNA editing sites identified at five different developmental stages were mutually confirmed. Finally, RNA editing frequency of one specific site was quantified by calculating the proportion of the nucleotide in reads with a substitution after mapping to the corresponding DNA template over the total reads in the same position. OrganellarGenomeDRAW [72] and Circos-0.69-5 [73] were used to visualize RNA editing sites and the corresponding RNA-seq coverage in mitochondria. In addition, RNA editing sites in *cox1*, *nad2*, *nad7* and *rpl2* were corrected by annotations from NCBI RefSeq under accession number DQ167400.1 because of incorrect annotations for these four genes under NC_011033.1.

### Detection of RNA editing variabilities during rice endosperm development

The *τ* value [74], originally developed for the identification of housekeeping and tissue-specific genes, was adopted in this study to estimate the RNA editing variability of any given site across different developmental stages of rice endosperm. For a given site, smaller *τ* value reflects relative constant editing frequencies across different conditions and larger *τ* value signifies variable editing frequencies.

### Correlation between editing frequency and evolutionary conservation of the corresponding recoded amino acids

To investigate the correlation between the editing efficiency of CDS-recoding sites and conservation of the corresponding recoded amino acids, alterations of amino acids caused by editing were obtained from mitochondrial genes, chromosomes, and genomes from accessions among land plants in RefSeq [75] and GenBank [76]. In addition, the correlation in *orf183* and *orf288* was filtered due to few accessions containing RNA editing sites in these two genes. Amino acids recoded by *ccmFn*-37, *ccmFn*-38, *mat-r*-295, *mat-r*-296, *rps19*-163, and *rps19*-164 were filtered because of more than one editing site in these corresponding codons. Multiple sequence alignment was conducted by ClustalW [77] in MEGA11 [78] and was used to calculate the percentage and conservation of amino acids at positions corresponding to RNA editing sites. In addition, amino acids at positions corresponding to RNA editing sites that exist in more than 80% species were kept because of gaps in the multiple sequence alignment. Furthermore, the correlation between editing frequency and conservation of amino acids was analyzed by Spearman’s correlation, fitted by the generalized linear model (GLM), and visualized by the R package ggpubr [79].

### RNA secondary structure prediction

To functionally investigate editing sites in intronic regions of mitochondrial genomes, editing sites in domain 5 (D5) in introns were selected, since D5 is the only domain in Group II introns that is sequence-conserved and can be identified by sequence alignment [80]. The secondary structure of *P.li.LSUI2* intron from the brown algae *Pylaiella littoralis* was revealed [81]. Next, D5 in introns of mitochondrial genes was searched by aligning with D5 of *P.li.LSUI2,* and D5 containing editing sites was kept. The secondary structure of D5 before and after RNA editing was predicted at 30 degree Celsius by MFold 2.3 (http://www.unafold.org/Dinamelt/applications/quickfold.php) [82].

### Mapping CDS-recoding sites to protein 3D structures

To investigate locations of CDS-recoding sites on 3D structures of proteins, experimentally verified protein 3D structures in Protein Data Bank (PDB) (https://www.rcsb.org) [83] were used with priority. Protein sequences in rice mitochondria and plastids were aligned with those in other plants with 3D annotations in PDB, and CDS-recoding sites were mapped to the 3D structures. Given that some annotations of 3D structures of proteins obtained by electron microscopy are not available in PDB, mapping of CDS-recoding sites to protein 3D structures was derived from annotations in Uniprot (https://www.uniprot.org) [84] and AlphaFold Protein Structure Database (AlphaFold DB, https://alphafold.ebi.ac.uk) [85].

### Protein 3D structure prediction and alignment

To investigate effects of CDS-recoding editing on 3D structures of rice mitochondrial proteins, ColabFold, the Alphafold2 server (https://colab.research.google.com/github/sokrypton/ColabFold) [86] was used to predict 3D structures of each protein before and after RNA editing, and hypothetical 3D structures were aligned and visualized by SuperPose (http://superpose.wishartlab.com) [87]. In addition, Root Mean Square Deviation (RMSD) greater than 2.5Å was considered a difference of protein 3D structures [46].

### Identification of PPR genes and prediction of their subcellular localization

To find all PPR genes in rice genome, PPRFinder (https://ppr.plantenergy.uwa.edu.au) [88, 89] was used to search PLS-class PPR genes and motifs, and PPRCODE (https://colab.research.google.com/github/YaoYinYing/PPRCODE_Guideline/

blob/master/PPRCODE.ipynb) [23], with built-in ScanProsite (https://prosite.expasy.org/scanprosite) [90], was used to recognize P-class PPR genes. Both PLS-zand P-class PPR codes were predicted by PPRCODE. Motifs and PPR codes of P-class PPR genes were revised by ScanProsite. In addition, motif gaps in PLS-class PPR genes were filled with inferred motifs as previously described [88], and ‘##’ is taken as the pseudo-PPR code of inferred motifs. PPR proteins with fewer than six motifs may include mis-annotated sequences or pseudogenes [88], and are filtered. Furthermore, subcellular localizations of PPR proteins were predicted by TargetP 2.0 [91], WoLF PSORT (https://wolfpsort.hgc.jp) [92], iPSORT (https://ipsort.hgc.jp) [93], Plant-mPLoc (http://www.csbio.sjtu.edu.cn/bioinf/plant-multi) [94], Predotar V1.04 (https://urgi.versailles.inra.fr/predotar) [95] and previously reported results [96]. For each PPR protein, the top 10% predictions generated by TargetP 2.0 and WoLF PSORT, and the top 20% predictions generated by Predotar V1.04 were retained, respectively. Only those predictions with the same subcellular localizations implemented by at least two aforementioned tools were kept for subsequent analysis.

### Prediction of PPR-RNA binding

To predict organelle RNA sequences bound by P-class PPRs, binding affinities of 62 PPR codes to RNA bases delineated by Yan et al [23] were utilized. Every RNA subsequence in each organelle gene was extracted and iterated, and the number of RNA bases was equal to P-class motifs. Then the binding of P-class PPRs to the RNA subsequence was scored. Specifically, the score of PPR-RNA binding (*S*) was estimated by

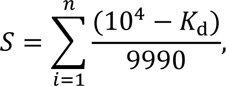

where *n* is the number of the subsequence bases or P-class PPR motifs and *K*_d_ (10 nm <= *K*_d_ <= 10^4^ nm; the smaller value of *K*_d_ means stronger binding affinity) is used to evaluate the binding affinity of each PPR code for one RNA base by isothermal titration calorimetry (ITC) experiments as previously described [23]. The prediction of P-class PPR proteins with at least six delineated PPR codes was retained. In addition, the prediction with the average binding score of each PPR code and RNA base less than 0.8 and the total binding score less than 10 would be discarded. Considering various motifs in PLS-class PPRs, the binding of PLS-class PPR-RNA was calculated as detailed in one previous study [97]. In addition, the repeats of PLS-class proteins bind the RNA subsequence that are from the 5’ terminal base to the 4^th^ base upstream of each RNA editing site [98].

## Supporting Information

**Figure S1. Density of distances to neighboring C-to-U editing sites in mitochondria.**

**Figure S2. Distribution of amino acids produced by CDS-recoding editing events on 3D structure of mitochondrial and plastid proteins.**

**Figure S3. 3D structure alignment of mitochondrial proteins before and after RNA editing.**

**Figure S4. The secondary structure of the domain 5 (D5) of the third and the only intron of *nad7* and *ccmFc* before (A, C) and after (B, D) editing.**

**Figure S5. RNA editing events in CDS (CDS-recoding and CDS-synonymous), intronic, intergenic and pseudogenic regions of rice mitochondrial genes at 3, 6, 9, 12 and 15 DAF.**

**Figure S6. The distribution and density of percentage of amino acids produced by RNA editing among land plants.**

**Figure S7. The distribution and density of editing frequency of CDS- recoding sites in rice endosperm mitochondria.**

**Figure S8. Correlation of the RNA editing frequency and evolutionary conservation of the recoded amino acid in rice mitochondria.**

**Figure S9. The correlation between conservation of amino acids and editing frequency of CDS-recoding sites in (A) liverworts, (B) mosses, (C) lycophytes, (D) seed plants, (E) ferns and (F) hornworts.**

**Figure S10. The correlation between conservation of the amino acids and editing frequency of CDS-recoding sites genes encoding cytochrome *c* biogenesis proteins among land plants.**

**Table S1. RNA editing events in mitochondria and plastids during rice endosperm development.**

**Table S2. Statistics of different types of RNA editing sites in mitochondria and plastids.**

**Table S3. RNA editing effect and codon position.**

**Table S4. Codon alterations caused by editing events in CDS regions.**

**Table S5 RNA editing events related to endosperm development in rice and other plants.**

**Table S6. Features of 214 C-to-U CDS-recoding sites in mitochondria during rice development.**

**Table S7. CDS-recoding sites on protein structures. Table S8. Species containing RNA editing sites.**

**Table S9. Multiple sequence alignment of 23 mitochondrial protein-coding genes after RNA editing.**

**Table S10. PPR genes in rice genome.**

**Table S11. Subcellular localization of PPR proteins in rice genome.**

**Table S12. PLS-type PPR protein binding to mitochondrial and plastid RNA.**

**Table S13. P-type PPR protein binding to mitochondrial and plastid RNA.**

**Table S14. RNA editing factor candidates of each C-to-U RNA editing site in mitochondria and plastids.**

**Table S15. Expression of PPR genes during rice endosperm development (at 3, 6, 9, 12 and 15 DAF).**

## Accession Numbers

The raw re-sequencing genome of rice and RNA-seq data of rice endosperm at 3, 6, 9, 12, and 15 DAF were deposited in the Genome Sequence Archive (GSA: CRA008750) [63] in the National Genomics Data Center, China National Center for Bioinformation / Beijing Institute of Genomics, Chinese Academy of Sciences [64]. A list of 254 mitochondrial genomes and genes of 122 land plants was obtained from NCBI RefSeq [75] and GenBank [76] and summarized in Table S8.

## Acknowledgments

These authors would like to thank Lili Hao, Shuhui Song, Lina Ma, Shuai Jiang, Dong Zou, Lin Liu, Qianpeng Li, Guangyi Niu, Zhao Li, Xiaonan Liu, Yang Zhang, Tongtong Zhu, Xing Zheng, Rong Pan, Sicheng Luo, Wei Jing, Yue Qi, Xinyu Zhou, Yuxin Qin, Wenzhuo Cheng, Zhixiang Yuan and Zhenxian Han for discussions and advice. This work was supported by grants from National Natural Science Foundation of China [32030021] and International Partnership Program of the Chinese Academy of Sciences [153F11KYSB20160008].

## Author Contributions

Z.Z., S.Y.W., and S.N.H. conceived, initiated and supervised this study. M.C. and L.X. wrote the manuscript. M.C., L.X., S.Y.W., X.Y.T., S.H.G., S.W., M.L.,

Y.S.Z., T.Y.X. and Y.C.C. analyzed the data. M.C., Y.Y.C, and Y.C. implemented the data curation. S.Y.W. provided the rice endosperm samples.

## References

1. Zhao M, Lin Y, Chen H. Improving nutritional quality of rice for human health. Theoretical and Applied Genetics. 2020;133(5):1397–413.

2. Chen H, He H, Zhou F, Yu H, Deng XW. Development of genomics-based genotyping platforms and their applications in rice breeding. Current opinion in plant biology. 2013;16(2):247–54.

3. Zhou S, Zhu S, Cui S, Hou H, Wu H, Hao B, et al. Transcriptional and post-transcriptional regulation of heading date in rice. New Phytologist. 2021;230(3):943–56.

4. An L, Tao Y, Chen H, He M, Xiao F, Li G, et al. Embryo-endosperm interaction and its agronomic relevance to rice quality. Frontiers in Plant Science. 2020;11:587641.

5. Zhou S-R, Yin L-L, Xue H-W. Functional genomics based understanding of rice endosperm development. Current opinion in plant biology. 2013;16(2):236–46.

6. Hoshikawa K. Anthesis, fertilization and development of caryopsis. Morphology. 1993.

7. Matsui T, Kobayasi K, Yoshimoto M, Hasegawa T, Tian X. Dependence of pollination and fertilization in rice (Oryza sativa L.) on floret height within the canopy. Field Crops Research. 2020;249:107741.

8. Wu X, Liu J, Li D, Liu CM. Rice caryopsis development II: Dynamic changes in the endosperm. Journal of Integrative Plant Biology. 2016;58(9):786–98.

9. Domínguez F, Cejudo FJ. Programmed cell death (PCD): an essential process of cereal seed development and germination. Frontiers in plant science. 2014;5:366.

10. Farooq MA, Zhang X, Zafar MM, Ma W, Zhao J. Roles of reactive oxygen species and mitochondria in seed germination. Frontiers in plant science. 2021;12:781734.

11. Møller IM, Rasmusson AG, Van Aken O. Plant mitochondria–past, present and future. The Plant Journal. 2021;108(4):912–59.

12. Notsu Y, Masood S, Nishikawa T, Kubo N, Akiduki G, Nakazono M, et al. The complete sequence of the rice (Oryza sativa L.) mitochondrial genome: frequent DNA sequence acquisition and loss during the evolution of flowering plants. Molecular Genetics and Genomics. 2002;268:434–45.

13. Wu J, Sun Y, Zhao Y, Zhang J, Luo L, Li M, et al. Deficient plastidic fatty acid synthesis triggers cell death by modulating mitochondrial reactive oxygen species. Cell research. 2015;25(5):621–33.

14. Blokhina O, Fagerstedt KV. Reactive oxygen species and nitric oxide in plant mitochondria: origin and redundant regulatory systems. Physiologia plantarum. 2010;138(4):447–62.

15. Giegé P, Grienenberger J, Bonnard G. Cytochrome c biogenesis in mitochondria. Mitochondrion. 2008;8(1):61–73.

16. Kim M, Lim J-H, Ahn CS, Park K, Kim GT, Kim WT, et al. Mitochondria-associated hexokinases play a role in the control of programmed cell death in Nicotiana benthamiana. The Plant Cell. 2006;18(9):2341–55.

17. Elena-Real CA, González-Arzola K, Pérez-Mejías G, Díaz-Quintana A, Velázquez-Campoy A, Desvoyes B, et al. Proposed mechanism for regulation of H2O2-induced programmed cell death in plants by binding of cytochrome c to 14-3-3 proteins. The Plant Journal. 2021;106(1):74–85.

18. Sultan LD, Mileshina D, Grewe F, Rolle K, Abudraham S, Głodowicz P, et al. The reverse transcriptase/RNA maturase protein MatR is required for the splicing of various group II introns in Brassicaceae mitochondria. The Plant Cell. 2016;28(11):2805–29.

19. Brown GG, Colas des Francs-Small C, Ostersetzer-Biran O. Group II intron splicing factors in plant mitochondria. Frontiers in plant science. 2014;5:35.

20. Janska H, Kwasniak M. Mitoribosomal regulation of OXPHOS biogenesis in plants. Frontiers in plant science. 2014;5:79.

21. Ghifari AS, Saha S, Murcha MW. The biogenesis and regulation of the plant oxidative phosphorylation system. Plant Physiology. 2023;192(2):728.

22. Takenaka M, Zehrmann A, Verbitskiy D, Haertel B, Brennicke A. RNA editing in plants and its evolution. Annual review of genetics. 2013;47:335–52.

23. Yan J, Yao Y, Hong S, Yang Y, Shen C, Zhang Q, et al. Delineation of pentatricopeptide repeat codes for target RNA prediction. Nucleic acids research. 2019;47(7):3728–38.

24. Okuda K, Shoki H, Arai M, Shikanai T, Small I, Nakamura T. Quantitative analysis of motifs contributing to the interaction between PLS-subfamily members and their target RNA sequences in plastid RNA editing. The Plant Journal. 2014;80(5):870–82.

25. Small ID, Schallenberg-Rüdinger M, Takenaka M, Mireau H, Ostersetzer-Biran O. Plant organellar RNA editing: what 30 years of research has revealed. The Plant Journal. 2020;101(5):1040–56.

26. Ren RC, Lu X, Zhao YJ, Wei YM, Wang LL, Zhang L, et al. Pentatricopeptide repeat protein DEK40 is required for mitochondrial function and kernel development in maize. Journal of experimental botany. 2019;70(21):6163–79.

27. Sosso D, Mbelo S, Vernoud V, Gendrot G, Dedieu A, Chambrier P, et al. PPR2263, a DYW-subgroup pentatricopeptide repeat protein, is required for mitochondrial nad5 and cob transcript editing, mitochondrion biogenesis, and maize growth. The Plant Cell. 2012;24(2):676–91.

28. Dai D, Jin L, Huo Z, Yan S, Ma Z, Qi W, et al. Maize pentatricopeptide repeat protein DEK53 is required for mitochondrial RNA editing at multiple sites and seed development. Journal of Experimental Botany. 2020;71(20):6246–61.

29. Ding S, Liu X-Y, Wang H-C, Wang Y, Tang J-J, Yang Y-Z, et al. SMK6 mediates the C-to-U editing at multiple sites in maize mitochondria. Journal of plant physiology. 2019;240:152992.

30. Wang G, Zhong M, Shuai B, Song J, Zhang J, Han L, et al. E+ subgroup PPR protein defective kernel 36 is required for multiple mitochondrial transcripts editing and seed development in maize and Arabidopsis. New Phytologist. 2017;214(4):1563–78.

31. Zang J, Zhang T, Zhang Z, Liu J, Chen H. DEFECTIVE KERNEL 56 functions in mitochondrial RNA editing and maize seed development. Plant Physiology. 2023:kiad598.

32. Wang Y, Liu X-Y, Yang Y-Z, Huang J, Sun F, Lin J, et al. Empty Pericarp21 encodes a novel PPR-DYW protein that is required for mitochondrial RNA editing at multiple sites, complexes I and V biogenesis, and seed development in maize. PLoS genetics. 2019;15(8):e1008305.

33. Sun F, Wang X, Bonnard G, Shen Y, Xiu Z, Li X, et al. Empty pericarp7 encodes a mitochondrial E-subgroup pentatricopeptide repeat protein that is required for ccmFN editing, mitochondrial function and seed development in maize. The Plant Journal. 2015;84(2):283–95.

34. Yang YZ, Ding S, Wang HC, Sun F, Huang WL, Song S, et al. The pentatricopeptide repeat protein EMP9 is required for mitochondrial ccmB and rps4 transcript editing, mitochondrial complex biogenesis and seed development in maize. New Phytologist. 2017;214(2):782–95.

35. Li XJ, Zhang YF, Hou M, Sun F, Shen Y, Xiu ZH, et al. Small kernel 1 encodes a pentatricopeptide repeat protein required for mitochondrial nad7 transcript editing and seed development in maize (Zea mays) and rice (Oryza sativa). The Plant Journal. 2014;79(5):797–809.

36. Liu Y-J, Xiu Z-H, Meeley R, Tan B-C. Empty pericarp5 encodes a pentatricopeptide repeat protein that is required for mitochondrial RNA editing and seed development in maize. The Plant Cell. 2013;25(3):868–83.

37. Kim SR, Yang JI, Moon S, Ryu CH, An K, Kim KM, et al. Rice OGR1 encodes a pentatricopeptide repeat–DYW protein and is essential for RNA editing in mitochondria. The Plant Journal. 2009;59(5):738–49.

38. Yang H, Wang Y, Tian Y, Teng X, Lv Z, Lei J, et al. Rice FLOURY ENDOSPERM22, encoding a pentatricopeptide repeat protein, is involved in both mitochondrial RNA splicing and editing and is crucial for endosperm development. J Integr Plant Biol. 2022. doi: 10.1111/jipb.13402. PubMed PMID: 36333887.

39. Li M, Xia L, Zhang Y, Niu G, Li M, Wang P, et al. Plant editosome database: a curated database of RNA editosome in plants. Nucleic Acids Research. 2019;47(D1):D170–D4.

40. Wu CS, Chaw SM. Evolution of mitochondrial RNA editing in extant gymnosperms. The Plant Journal. 2022;111(6):1676–87.

41. Maldonado M, Abe KM, Letts JA. A structural perspective on the RNA editing of plant respiratory complexes. International Journal of Molecular Sciences. 2022;23(2):684.

42. Koo HJ, Yang T-J. RNA Editing may stabilize membrane-embedded proteins by increasing phydrophobicity: a study of Zanthoxylum piperitum and Z. schinifolium chloroplast NdhG. Gene. 2020;746:144638.

43. Wang Y, Li H, Huang Z-Q, Ma B, Yang Y-Z, Xiu Z-H, et al. Maize PPR-E proteins mediate RNA C-to-U editing in mitochondria by recruiting the trans deaminase PCW1. The Plant Cell. 2023;35(1):529–51.

44. Liu X-Y, Jiang R-C, Wang Y, Tang J-J, Sun F, Yang Y-Z, et al. ZmPPR26, a DYW-type pentatricopeptide repeat protein, is required for C-to-U RNA editing at atpA-1148 in maize chloroplasts. Journal of Experimental Botany. 2021;72(13):4809–21.

45. Yura K, Go M. Correlation between amino acid residues converted by RNA editing and functional residues in protein three-dimensional structures in plant organelles. BMC Plant Biology. 2008;8:1–11.

46. Bolhuis P. Sampling kinetic protein folding pathways using all-atom models. Computer Simulations in Condensed Matter Systems: From Materials to Chemical Biology Volume 1: Springer; 2006. p. 393–433.

47. Lang BF, Laforest M-J, Burger G. Mitochondrial introns: a critical view. Trends in Genetics. 2007;23(3):119–25.

48. Toor N, Keating KS, Fedorova O, Rajashankar K, Wang J, Pyle AM. Tertiary architecture of the Oceanobacillus iheyensis group II intron. RNA. 2010;16(1):57–69.

49. Xu C, Song S, Yang YZ, Lu F, Zhang MD, Sun F, et al. DEK46 performs C-to-U editing of a specific site in mitochondrial nad7 introns that is critical for intron splicing and seed development in maize. The Plant Journal. 2020;103(5):1767–82.

50. Ichinose M, Sugita M. RNA editing and its molecular mechanism in plant organelles. Genes. 2016;8(1):5.

51. Duan Y, Tang X, Lu J. Evolutionary driving forces of A-to-I editing in metazoans. Wiley Interdisciplinary Reviews: RNA. 2022;13(1):e1666.

52. Bi C, Lu N, Xu Y, He C, Lu Z. Characterization and analysis of the mitochondrial genome of common bean (Phaseolus vulgaris) by comparative genomic approaches. International journal of molecular sciences. 2020;21(11):3778.

53. Loiacono FV, Walther D, Seeger S, Thiele W, Gerlach I, Karcher D, et al. Emergence of Novel RNA-Editing Sites by Changes in the Binding Affinity of a Conserved PPR Protein. Molecular Biology and Evolution. 2022;39(12):msac222.

54. Wang W, Wu Y, Messing J. Genome-wide analysis of pentatricopeptide-repeat proteins of an aquatic plant. Planta. 2016;244:893–9.

55. Lurin C, Andreés C, Aubourg S, Bellaoui M, Bitton F, Bruyère C, et al. Genome-wide analysis of Arabidopsis pentatricopeptide repeat proteins reveals their essential role in organelle biogenesis. The Plant Cell. 2004;16(8):2089–103.

56. Sheahan T, Wieden H-J. Ribosomal protein S1 improves the protein yield of an in vitro reconstituted cell-free translation system. ACS Synthetic Biology. 2022;11(2):1004–8.

57. Fargo DC, Boynton JE, Gillham NW. Chloroplast ribosomal protein S7 of Chlamydomonas binds to chloroplast mRNA leader sequences and may be involved in translation initiation. The Plant Cell. 2001;13(1):207–18.

58. Cukras AR, Southworth DR, Brunelle JL, Culver GM, Green R. Ribosomal proteins S12 and S13 function as control elements for translocation of the mRNA: tRNA complex. Molecular cell. 2003;12(2):321–8.

59. Liu R, Cao S-K, Sayyed A, Yang H-H, Zhao J, Wang X, et al. The DYW-subgroup pentatricopeptide repeat protein PPR27 interacts with ZmMORF1 to facilitate mitochondrial RNA editing and seed development in maize. Journal of Experimental Botany. 2020;71(18):5495–505.

60. Ren RC, Yan XW, Zhao YJ, Wei YM, Lu X, Zang J, et al. The novel E-subgroup pentatricopeptide repeat protein DEK55 is responsible for RNA editing at multiple sites and for the splicing of nad1 and nad4 in maize. BMC Plant Biology. 2020;20(1):1–15.

61. Takatsuka A, Kazama T, Toriyama K. Cytoplasmic male sterility-associated mitochondrial gene orf312 derived from rice (Oryza sativa L.) cultivar Tadukan. Rice. 2021;14(1):1–11.

62. Edera AA, Gandini CL, Sanchez-Puerta MV. Towards a comprehensive picture of C-to-U RNA editing sites in angiosperm mitochondria. Plant Molecular Biology. 2018;97:215–31.

63. Chen T, Chen X, Zhang S, Zhu J, Tang B, Wang A, et al. The genome sequence archive family: toward explosive data growth and diverse data types. Genomics, Proteomics & Bioinformatics. 2021;19(4):578–83.

64. CNCB-NGDC Members and Partners. Database Resources of the National Genomics Data Center, China National Center for Bioinformation in 2024. Nucleic Acids Research. 2024;52(D1):D18–D32.

65. Bolger AM, Lohse M, Usadel B. Trimmomatic: a flexible trimmer for Illumina sequence data. Bioinformatics. 2014;30(15):2114–20. doi: 10.1093/bioinformatics/btu170. PubMed PMID: 24695404; PubMed Central PMCID: PMCPMC4103590.

66. Notsu Y, Masood S, Nishikawa T, Kubo N, Akiduki G, Nakazono M, et al. The complete sequence of the rice (Oryza sativa L.) mitochondrial genome: frequent DNA sequence acquisition and loss during the evolution of flowering plants. Molecular Genetics and Genomics. 2002;268(4):434–45.

67. Hiratsuka J, Shimada H, Whittier R, Ishibashi T, Sakamoto M, Mori M, et al. The complete sequence of the rice (Oryza sativa) chloroplast genome: intermolecular recombination between distinct tRNA genes accounts for a major plastid DNA inversion during the evolution of the cereals. Mol Gen Genet. 1989;217(2-3):185–94. PubMed PMID: 2770692.

68. Kawahara Y, de la Bastide M, Hamilton JP, Kanamori H, McCombie WR, Ouyang S, et al. Improvement of the Oryza sativa Nipponbare reference genome using next generation sequence and optical map data. Rice. 2013;6(1):1–10.

69. Kim D, Langmead B, Salzberg SL. HISAT: a fast spliced aligner with low memory requirements. Nat Methods. 2015;12(4):357–60. doi: 10.1038/nmeth.3317. PubMed PMID: 25751142; PubMed Central PMCID: PMCPMC4655817.

70. Pertea M, Pertea GM, Antonescu CM, Chang TC, Mendell JT, Salzberg SL. StringTie enables improved reconstruction of a transcriptome from RNA-seq reads. Nat Biotechnol. 2015;33(3):290–5. doi: 10.1038/nbt.3122. PubMed PMID: 25690850; PubMed Central PMCID: PMCPMC4643835.

71. Picardi E, Pesole G. REDItools: high-throughput RNA editing detection made easy. Bioinformatics. 2013;29(14):1813–4. doi: 10.1093/bioinformatics/btt287. PubMed PMID: 23742983.

72. Lohse M, Drechsel O, Kahlau S, Bock R. OrganellarGenomeDRAW--a suite of tools for generating physical maps of plastid and mitochondrial genomes and visualizing expression data sets. Nucleic Acids Res. 2013;41(Web Server issue):W575–81. doi: 10.1093/nar/gkt289. PubMed PMID: 23609545; PubMed Central PMCID: PMCPMC3692101.

73. Krzywinski M, Schein J, Birol I, Connors J, Gascoyne R, Horsman D, et al. Circos: an information aesthetic for comparative genomics. Genome Res. 2009;19(9):1639–45. doi: 10.1101/gr.092759.109. PubMed PMID: 19541911; PubMed Central PMCID: PMCPMC2752132.

74. Yanai I, Benjamin H, Shmoish M, Chalifa-Caspi V, Shklar M, Ophir R, et al. Genome-wide midrange transcription profiles reveal expression level relationships in human tissue specification. Bioinformatics. 2005;21(5):650–9. doi: 10.1093/bioinformatics/bti042. PubMed PMID: 15388519.

75. O’Leary NA, Wright MW, Brister JR, Ciufo S, Haddad D, McVeigh R, et al. Reference sequence (RefSeq) database at NCBI: current status, taxonomic expansion, and functional annotation. Nucleic acids research. 2016;44(D1):D733–D45.

76. Benson DA, Cavanaugh M, Clark K, Karsch-Mizrachi I, Lipman DJ, Ostell J, et al. GenBank. Nucleic acids research. 2012;41(D1):D36–D42.

77. Thompson JD, Higgins DG, Gibson TJ. CLUSTAL W: improving the sensitivity of progressive multiple sequence alignment through sequence weighting, position-specific gap penalties and weight matrix choice. Nucleic acids research. 1994;22(22):4673–80.

78. Tamura K, Stecher G, Kumar S. MEGA11: molecular evolutionary genetics analysis version 11. Molecular biology and evolution. 2021;38(7):3022–7.

79. Kassambara A. ggpubr:“ggplot2” based publication ready plots. R package version 04 0. 2020;438.

80. Bonen L, Vogel J. The ins and outs of group II introns. TRENDS in Genetics. 2001;17(6):322–31.

81. Robart AR, Chan RT, Peters JK, Rajashankar KR, Toor N. Crystal structure of a eukaryotic group II intron lariat. Nature. 2014;514(7521):193-7.

82. Zuker M. Mfold web server for nucleic acid folding and hybridization prediction. Nucleic acids research. 2003;31(13):3406–15.

83. Berman HM, Westbrook J, Feng Z, Gilliland G, Bhat TN, Weissig H, et al. The protein data bank. Nucleic acids research. 2000;28(1):235–42.

84. UniProt: the Universal Protein knowledgebase in 2023. Nucleic Acids Research. 2023;51(D1):D523–D31.

85. Varadi M, Anyango S, Deshpande M, Nair S, Natassia C, Yordanova G, et al. AlphaFold Protein Structure Database: massively expanding the structural coverage of protein-sequence space with high-accuracy models. Nucleic acids research. 2022;50(D1):D439–D44.

86. Mirdita M, Schütze K, Moriwaki Y, Heo L, Ovchinnikov S, Steinegger M. ColabFold: making protein folding accessible to all. Nature methods. 2022;19(6):679–82.

87. Maiti R, Van Domselaar GH, Zhang H, Wishart DS. SuperPose: a simple server for sophisticated structural superposition. Nucleic acids research. 2004;32(suppl_2):W590–W4.

88. Cheng S, Gutmann B, Zhong X, Ye Y, Fisher MF, Bai F, et al. Redefining the structural motifs that determine RNA binding and RNA editing by pentatricopeptide repeat proteins in land plants. The Plant Journal. 2016;85(4):532–47.

89. Gutmann B, Royan S, Schallenberg-Rüdinger M, Lenz H, Castleden IR, McDowell R, et al. The expansion and diversification of pentatricopeptide repeat RNA-editing factors in plants. Molecular plant. 2020;13(2):215–30.

90. De Castro E, Sigrist CJ, Gattiker A, Bulliard V, Langendijk-Genevaux PS, Gasteiger E, et al. ScanProsite: detection of PROSITE signature matches and ProRule-associated functional and structural residues in proteins. Nucleic acids research. 2006;34(suppl_2):W362–W5.

91. Armenteros JJA, Salvatore M, Emanuelsson O, Winther O, Von Heijne G, Elofsson A, et al. Detecting sequence signals in targeting peptides using deep learning. Life science alliance. 2019;2(5).

92. Horton P, Park K-J, Obayashi T, Fujita N, Harada H, Adams-Collier C, et al. WoLF PSORT: protein localization predictor. Nucleic acids research. 2007;35(suppl_2):W585–W7.

93. Bannai H, Tamada Y, Maruyama O, Nakai K, Miyano S. Extensive feature detection of N-terminal protein sorting signals. Bioinformatics. 2002;18(2):298–305.

94. Chou K-C, Shen H-B. Plant-mPLoc: a top-down strategy to augment the power for predicting plant protein subcellular localization. PloS one. 2010;5(6):e11335.

95. Small I, Peeters N, Legeai F, Lurin C. Predotar: a tool for rapidly screening proteomes for N-terminal targeting sequences. Proteomics. 2004;4(6):1581–90.

96. Chen G, Zou Y, Hu J, Ding Y. Genome-wide analysis of the rice PPR gene family and their expression profiles under different stress treatments. BMC genomics. 2018;19(1):1–14.

97. Gutmann B, Millman M, Sanglard LVP, Small I, Francs-Small CCd. The Pentatricopeptide Repeat Protein MEF100 Is Required for the Editing of Four Mitochondrial Editing Sites in Arabidopsis. Cells. 2021;10(2):468.

98. Takenaka M, Zehrmann A, Brennicke A, Graichen K. Improved computational target site prediction for pentatricopeptide repeat RNA editing factors. PloS one. 2013;8(6):e65343.

